# Molecular chaperones accelerate the evolution of their protein clients in yeast

**DOI:** 10.1101/552349

**Authors:** David Alvarez-Ponce, José Aguilar-Rodríguez, Mario A. Fares

**Author notes:** These authors contributed equally to this work (David Alvarez-Ponce, José Aguilar-Rodríguez). Posthumous author (Mario A. Fares). **Corresponding author:** David Alvarez-Ponce, Department of Biology, University of Nevada, Reno. 1664 N. Virginia Street, Reno, NV 89557.

## Abstract

Protein stability is a major constraint on protein evolution. Molecular chaperones, also known as heat-shock proteins, can relax this constraint and promote protein evolution by diminishing the deleterious effect of mutations on protein stability and folding. This effect, however, has only been stablished for a few chaperones. Here, we use a comprehensive chaperone-protein interaction network to study the effect of all yeast chaperones on the evolution of their protein substrates, that is, their clients. In particular, we analyze how yeast chaperones affect the evolutionary rates of their clients at two very different evolutionary time scales. We first study the effect of chaperone-mediated folding on protein evolution over the evolutionary divergence of *Saccharomyces cerevisiae* and *S. paradoxus*. We then test whether yeast chaperones have left a similar signature on the patterns of standing genetic variation found in modern wild and domesticated strains of *S. cerevisiae*. We find that genes encoding chaperone clients have diverged faster than genes encoding nonclient proteins when controlling for their number of protein-protein interactions. We also find that genes encoding client proteins have accumulated more intra-specific genetic diversity than those encoding nonclient proteins. In a number of multivariate analyses, controlling by other well-known factors that affect protein evolution, we find that chaperone dependence explains the largest fraction of the observed variance in the rate of evolution at both evolutionary time scales. Chaperones affecting rates of protein evolution mostly belong to two major chaperone families: Hsp70s and Hsp90s. Our analyses show that protein chaperones, by virtue of their ability to buffer destabilizing mutations and their role in modulating protein genotype-phenotype maps, have a considerable accelerating effect on protein evolution.

## INTRODUCTION

Proteins within the proteome of any organism evolve at very different rates: whereas some proteins remain largely unaltered during long evolutionary periods, others can undergo fast evolutionary changes (Zuckerkandl and Pauling 1965; Zuckerkandl 1976; Li, et al. 1985). The reasons for this diversity in rates of protein evolution are still a subject of intense debate (Rocha 2006; Alvarez-Ponce 2014; Zhang and Yang 2015). A number of factors have been shown to affect rates of evolution, including gene expression levels (Pál, et al. 2001; Drummond, et al. 2005), expression breadth in multicellular organisms (Duret and Mouchiroud 2000; Wright, et al. 2004; Zhang and Li 2004; Alvarez-Ponce and Fares 2012), essentiality (Hurst and Smith 1999; Jordan, et al. 2002; Alvarez-Ponce, et al. 2016; Aguilar-Rodríguez and Wagner 2018), duplicability (Nembaware, et al. 2002; Yang, et al. 2003; Pegueroles, et al. 2013) and the number of protein–protein interactions (Fraser, et al. 2002; Hahn and Kern 2005; Alvarez-Ponce and Fares 2012). However, a comprehensive understanding of which factors affect rates of protein evolution, their relative impacts on rates of evolution, and the molecular mechanisms underlying these impacts, is lacking.

Molecular chaperones (Ellis 1987) help other proteins achieve their functional and three-dimensional native conformations, prevent protein aggregation, and restore the native conformation of proteins destabilized by environmental perturbations (Hartl and Hayer-Hartl 2009; Hartl, et al. 2011). As such, they can render neutral certain amino acid substitutions that would otherwise (in the absence of chaperones) be deleterious (or at least diminish their negative fitness effects) (Tokuriki and Tawfik 2009). Chaperones thus represent an extrinsic source of protein robustness: They can increase the tolerance of a protein phenotype (e.g., protein structure responsible for the protein function) against mutational insults. Therefore, chaperones can be not only a source of environmental robustness, but also of mutational robustness (Jarosz, et al. 2010; Lauring, et al. 2013; Fares 2015; Payne and Wagner 2018). That is, chaperones can effectively buffer certain types of mutations in proteins, and thus are expected to contribute to the accumulation of genetic variation, and to increase the rates of evolution of their clients.

This increased rate of protein evolution of the clients of certain chaperones has been detected at the genomic level in a number of studies. Comparative analysis of bacterial genomes shows that the GroEL/ES chaperonin system can increase the evolutionary rate of its client proteins: after controlling for confounding factors, proteins that are clients of the system evolve faster on average than those that are not clients (Bogumil & Dagan 2010; Williams & Fares 2010). The bacterial DnaK also accelerates the rate of evolution of its clients (Aguilar-Rodríguez et al. 2016; Kadibalban et al. 2016). In yeast, Hsp90 clients evolve faster than their nonclient paralogs (Lachowiec et al. 2013), and distinct groups of proteins interacting with different chaperones evolve at different rates (Bogumil et al. 2012). In mammals, kinases with higher binding affinity to Hsp90 evolve faster than kinases with lower binding affinity (Lachowiec et al. 2015). It has also been shown that both co- and post-translationally acting chaperones can promote non-conservative amino acid substitutions, more likely destabilizing mutations, in their clients (Pechmann and Frydman 2014).

However, most studies so far have focused on individual chaperones and lineages, and the effect of most chaperones on protein evolution remains unknown. In this study, we evaluate the effect of all yeast protein chaperones on the evolution of their protein clients. We conducted a comprehensive analysis of the chaperone–client interaction network of 35 chaperones in yeast (Gong, et al. 2009). This network was established with TAP-tag pulldown assays followed by both liquid chromatography tandem mass spectrometry (LC-MS/MS) and by matrix-assisted laser desorption/ionization-time of flight mass spectrometry (MALDI-TOF). We used this high-quality network to evaluate whether chaperone clients evolve faster in yeast, and also to measure the contribution of different chaperone families to this acceleration of the rate of evolution. We show that many chaperones accelerate not only the rates of evolution of their clients, but also their levels of nonsynonymous polymorphism.

## RESULTS

### Yeast chaperone clients evolve slower than non-clients

We classified all *Saccharomyces cerevisiae* proteins into three classes: chaperones (*n* = 35), co-chaperones (*n* = 29), and others (*n* = 6653), using the chaperone and co-chaperone list by Gong et al. (2009). The latter class was further classified into chaperone clients (those that interact with any of the chaperones according to the dataset of Gong et al. (2009), *n* = 4209) and non-clients (all remaining proteins*, n* = 2444).

For each *S. cerevisiae* gene, the most likely ortholog in *Saccharomyces paradoxus* was identified using a best-reciprocal-hit approach (see *Methods*), and the rate of protein evolution was measured from the nonsynonymous to synonymous divergence ratio (*d*_N_/*d*_S_). These species diverged from a common ancestor ∼5–10 million years ago (Dori-Bachash, et al. 2011). Orthologs could be identified for 5603 of the *S. cerevisiae* genes. Values of *d*_N_/*d*_S_ above 8 were removed, as they probably represent artifacts (10 genes were removed). The mean *d*_N_/*d*_S_ value was 0.1553, and the median was 0.0970, consistent with prior results (e.g., Alvarez-Ponce, et al. 2017). After applying these filters, a total of 3958 clients and 1574 non-clients were available for analysis. All remaining genes were excluded from further analyses.

Clients exhibit substantially lower *d*_N_/*d*_S_ values (median: 0.0930) than non-clients (median: 0.1149; Mann–Whitney’s *U* test, *P*-value = 9.48×10^−22^; Fig. 1; Table 1). They also exhibit lower *d*_N_ and higher *d*_S_ values (Fig. 1; Table 1). Next, we considered whether the number of chaperones of which each protein is client correlates with its rate of evolution. Among the 3958 genes that have an ortholog in *S. paradoxus* and are clients of at least one chaperone, *d*_N_/*d*_S_ negatively correlates with the number of chaperones (Spearman’s rank correlation coefficient, ρ = −0.0784, *P* = 7.79×10^−7^). The number of chaperones also correlates with *d*_N_ (ρ = −0.0596, *P* = 0.0002) and, to a lesser extent, with *d*_S_ (ρ = 0.0323, *P* = 0.0422).

**Table 1.** Comparison of yeast chaperone clients vs. non-clients

|  | Chaperone clients |  |  | Non-clients |  |  | *P*-value |
| --- | --- | --- | --- | --- | --- | --- | --- |
|  | *n* | Mean | Median | *n* | Mean | Median |  |
| $d_N/d_S$ | 3958 | 0.1165 | 0.0930 | 1574 | **0.2563** | **0.1149** | $9.48 \times 10^{-22}***$ |
| $d_N$ | 3958 | 0.0432 | 0.0355 | 1574 | **0.0653** | **0.0411** | $5.66 \times 10^{-12}***$ |
| $d_S$ | 3958 | **0.3795** | **0.3817** | 1574 | 0.3722 | 0.3655 | $2.80 \times 10^{-11}***$ |
| Number of protein–protein interactions | 3875 | **30.3130** | **16** | 1265 | 18.7107 | 8 | $3.62 \times 10^{-53}***$ |
| Expression level | 3434 | **71.1133** | **23** | 1184 | 69.5845 | 20 | $1.71 \times 10^{-5}***$ |
| Protein length | 3958 | **553.5682** | **462** | 1574 | 327.5172 | 269 | $3.10 \times 10^{-138}***$ |

For each pair of clients vs. non-client values, the highest value is shown in bold face. *P*-values correspond to the Mann–Whitney test. Significance levels: \**P* < 0.05, \*\**P* < 0.001, \*\*\**P* < 10<sup>−5</sup>.

**Fig. 1.**
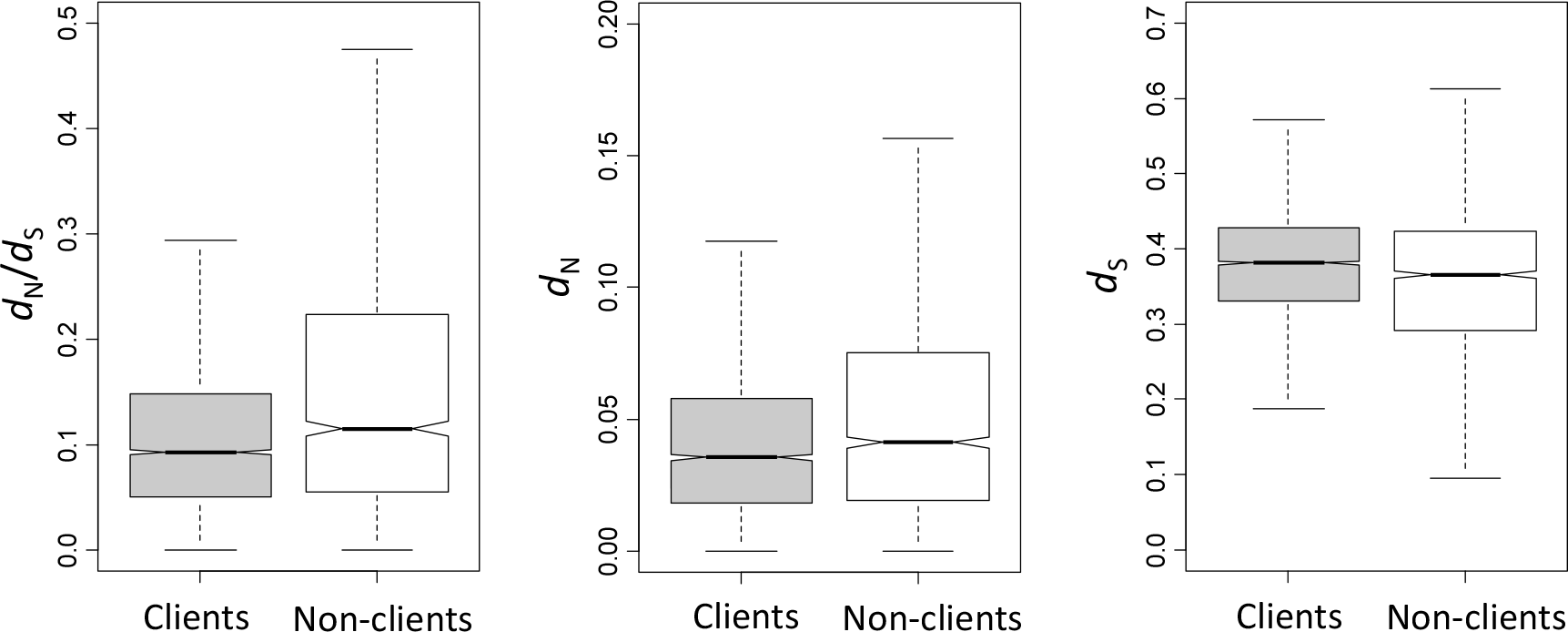
Rates of evolution of yeast chaperone clients and non-clients. Outliers (those above the 90^th^ and below the 10^th^ percentiles) are not shown.

We next considered whether chaperone clients may be enriched in proteins encoded by genes under positive selection. For each *S. cerevisiae* gene, we identified its most likely orthologs in another 4 species of the genus *Saccharomyces* (*S. paradoxus*, *S. mikatae*, *S. kudriavzevii*, and *S. bayanus*). Only genes with a putative ortholog in all species (*n* = 2047) were included in this analysis. The M8 vs. M7 test (Yang 2000) was used to identify signatures of positive selection (see *Methods*). Among chaperone clients, 19 genes (3.40%) were encoded by genes under positive selection. Among non-clients, 72 (4.84%) were encoded by genes with signatures of positive selection. The fraction of genes under positive selection was not significantly different between clients and non-clients (Fisher’s exact test, *P* = 0.0967).

### The low rates of evolution of chaperone clients is not due to their expression levels, essentiality or duplicability

Rates of protein evolution are affected by a number of parameters, including expression levels (Pál, et al. 2001; Drummond, et al. 2005), gene essentiality (Hurst and Smith 1999; Jordan, et al. 2002; Alvarez-Ponce, et al. 2016), gene duplicability (Nembaware, et al. 2002; Yang, et al. 2003; Pegueroles, et al. 2013) and number of protein–protein interactions (Fraser, et al. 2002; Hahn and Kern 2005; Alvarez-Ponce and Fares 2012) (for review, see Rocha 2006; Alvarez-Ponce 2014; Zhang and Yang 2015). Clients and non-clients differ in all these parameters (Table 1), and thus it is conceivable that the observed differences in the rates of evolution of clients and non-clients (Fig. 1; Table 1) might be a byproduct of differences in these factors. In order to discard this possibility, we conducted a number of controls.

Expression level seems to be a major determinant of protein’s rates of evolution, with highly expressed genes tending to be more selectively constrained (Pál, et al. 2001; Drummond, et al. 2005; Drummond, et al. 2006). In agreement with prior results, we observed a negative correlation between expression levels and *d*_N_/*d*_S_ (ρ = −0.4138, *P* = 1.73×10^−190^). Chaperone clients are more highly expressed than non-clients (median expression level for clients: 23; median expression level for non-clients: 20; Mann–Whitney test, *P* = 1.71×10^−5^). This raises the possibility that the lower rates of evolution of clients might be a byproduct of clients being more highly expressed. However, partial correlation analysis shows that the relationship between “chaperone dependence” (a dummy variable taking the value of 1 if the protein is client of at least one chaperone, and 0 otherwise) and *d*_N_/*d*_S_ is independent of expression level (partial Spearman’s rank correlation coefficient, ρ = −0.0414, *P* = 0.0049). Furthermore, among chaperone clients, the partial correlation between *d*_N_/*d*_S_ and number of chaperones controlling for expression level is significantly negative (ρ = −0.0643, *P* = 0.00016).

Proteins encoded by essential genes tend to be more constrained than those encoded by non-essential genes (Hurst and Smith 1999; Alvarez-Ponce, et al. 2016). Among the 3958 chaperone clients with *d*_N_/*d*_S_ information, 831 (i.e., 21%) are essential. Among the 1574 non-clients, only 228 (14.5%) are essential. Thus, clients are enriched in essential genes (Fisher exact test, *P* < 10^−6^), which could potentially explain their low evolutionary rates. To discard this possibility, we analyzed essential and non-essential genes separately, and in both cases clients exhibited a lower *d*_N_/*d*_S_. Among essential genes, the median *d*_N_/*d*_S_ was 0.0692 for clients and 0.0913 for non-clients (Mann–Whitney test, *P* = 0.0016). Among non-essential genes, the median *d*_N_/*d*_S_ was 0.0990 for clients and 0.1179 for non-clients (Mann–Whitney test, *P* = 2.06×10^−16^).

Proteins encoded by duplicated genes tend to evolve slower than those encoded by singleton genes (Nembaware, et al. 2002; Yang, et al. 2003), in spite of the fact that gene duplication transiently accelerates protein evolution (Han, et al. 2009; Pegueroles, et al. 2013). Among clients, 1684 (42.54%) are encoded by duplicated genes, and among non-clients, 547 (34.75%) are encoded by duplicated genes; i.e., clients are enriched in proteins encoded by duplicated genes (Fisher’s exact test, *P* < 10^−6^), which might account for their slow evolution. To discard this possibility, we analyzed singleton and duplicated genes separately. Among singletons, clients exhibit lower *d*_N_/*d*_S_ values (median = 0.1048) than non-clients (median = 0.1462; Mann–Whitney’s *U* test, *P* = 9.45×10^−27^). Among the less numerous duplicates, clients also exhibited lower *d*_N_/*d*_S_ values, but the differences were not significant (median for clients: 0.0752, median for non-clients: 0.0786, *P* = 0.3210). In addition, among clients, the number of chaperones significantly correlates with *d*_N_/*d*_S_, among both singletons (ρ = −0.0705, *P* = 0.0008) and duplicates (ρ = −0.0745, *P* = 0.0022). These results indicate that the lower rates of evolution of chaperone clients are not due to their enrichment in proteins encoded by duplicated genes.

### Controlling for number of physical interactions reveals that chaperone dependence accelerates protein evolution

The number of protein–protein interactions with which a protein interacts (degree centrality) negatively correlates with its rate of evolution (Fraser, et al. 2002; Hahn and Kern 2005; Alvarez-Ponce and Fares 2012), a pattern that was also apparent in our dataset (ρ = −0.2788, *P* = 2.14×10^−92^). This, together with the fact that chaperone clients tend to exhibit more protein– protein interactions (median = 16) than non-clients (median = 8; Mann–Whitney *U* test, *P* = 3.62×10^−53^), might account for the low rates of evolution of chaperone clients.

Indeed, the partial correlation between *d*_N_/*d*_S_ and chaperone dependence while controlling for degree is significantly positive (ρ = 0.0507, *P* = 0.0003), as is the partial correlation between the *d*_N_/*d*_S_ values of clients and their number of chaperones while controlling for degree (ρ = 0.0181, *P* = 2.71×10^−6^). These results indicate that chaperones accelerate the rates of evolution of their clients.

We repeated these analyses using degree values computed from a subset of protein– protein interactions of high quality (interactions identified either by low-throughput screens or by two or more high-throughput screens). This reduced the number of genes for which available network data was available from 5140 to 4011. The partial correlation between *d*_N_/*d*_S_ and chaperone dependence while controlling for degree remains significantly positive (ρ = 0.0405, *P* = 0.0104), while the correlation between *d*_N_/*d*_S_ and the number of chaperones with which clients interact was not significant (ρ = 0.0055, *P* = 0.7546).

To further validate our results, we binned proteins into 7 degree classes: 1–5 interactions (*n* = 1224), 6–10 interactions (*n* = 927), 11–15 interactions (*n* = 657), 16–20 interactions (*n* = 410), 21–25 interactions (*n* = 351), 26–30 interactions (*n* = 243), and >30 interactions (*n* = 1328). Within each of the classes, chaperone clients exhibited a lower median *d*_N_/*d*_S_ than non-clients (Fig. 2), with significant differences in the classes of degree 15–20 (one-tailed Mann–Whitney test, *P* = 4.30×10^−5^) and degree > 30 (*P* = 0.0385). In addition, the observation that in all 7 categories clients have a higher median *d*_N_/*d*_S_ is not expected at random (binomial test, *P* = 0.0156).

**Fig. 2.**
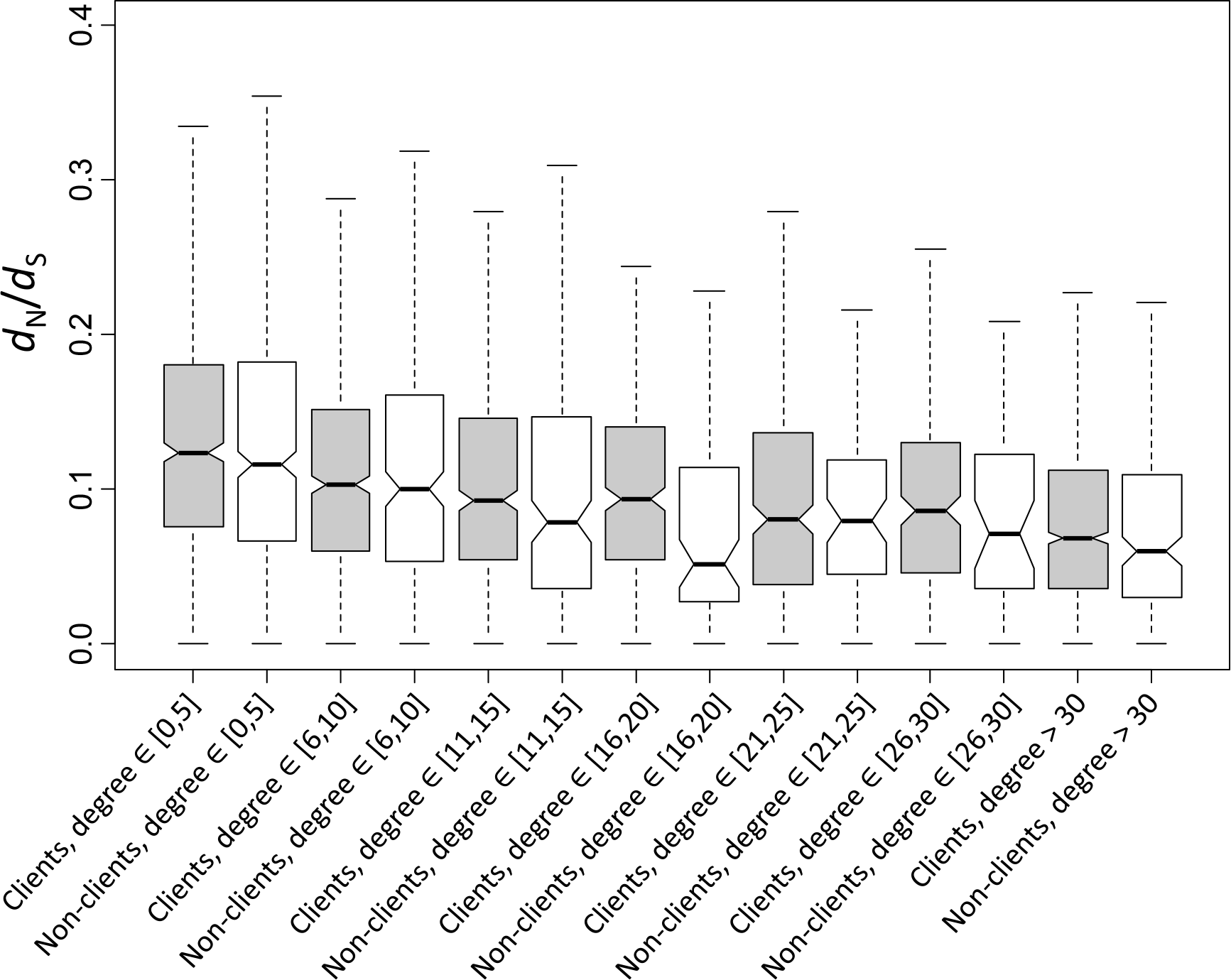
Comparison of the rate of evolution of clients and non-clients with different degrees. Clients are represented in grey and non-clients in white. Outliers (those above the 90^th^ and below the 10^th^ percentiles) are not shown.

### Multivariate analyses confirm the accelerating effect of chaperones on the evolution of their clients

We performed a multivariate regression analysis to study the relative influence of all the studied factors (chaperone dependence, expression level, number of protein–protein interactions, duplicability, and essentiality) simultaneously. We regressed *d*_N_*/d*_S_ against the five biological factors, and found that all make a significant contribution to the regression and that the overall *R*^*2*^ is 0.219 (Table 2). Chaperone dependence was the only factor with a positive coefficient, indicating that chaperone dependence increases protein evolutionary rates. Multivariate regression assumes that the predictor variables are statistically independent. To evaluate if our predictors intercorrelate we used the variance inflation factor (VIF) to quantify the degree of collinearity. We found a VIF of 1.28, which indicates that while collinearity is present in our model, it is rather low. Nevertheless, multivariate regression can produce spurious results in the presence of both collinearity and noise (Drummond, et al. 2006), and our variables are affected by noisy measurements. Therefore, we also performed a principal component regression analysis, which is an established method to study the relative contributions of different determinants of protein evolutionary rates (Drummond, et al. 2006), although it is not entirely insensitive to noise (Plotkin and Fraser 2007). Principal component regression finds new variables, called principal components, which are linear combinations of the original predictor variables, and then regresses the response variable against them. We performed principal component regression using the same predictor variables as above. Table 3 show numerical data from the analysis, while Fig. 3 shows these data graphically.

**Table 2.** Multiple linear regression of divergence data

| | $d_N/d_S$ | $d_N$ | $d_S$ |
| --- | --- | --- | --- |
| Chaperone dependence | 0.17*** | 0.14*** | 0.06*** |
| Number of protein–protein interactions | −0.16*** | −0.13*** | −0.01* |
| Expression level | −0.34*** | −0.31*** | −0.09*** |
| Duplicability | −0.38*** | −0.34*** | −0.02* |
| Essentiality | −0.35*** | −0.30*** | −0.01 |

Regression coefficients are shown. Significance levels: \* $P < 0.05$ , \*\* $P < 0.001$ , \*\*\* $P < 10^{-5}$ .

**Table 3.** Results from the principal component regression analysis of divergence data

|  | Principal components |  |  |  |  |  |
| --- | --- | --- | --- | --- | --- | --- |
|  | 1 | 2 | 3 | 4 | 5 | All |
| Percentage of explained variance in |  |  |  |  |  |  |
| $d_N/d_S$ | 13.09*** | 2.04*** | 6.11*** | 0.22*** | 0.39*** | 21.85 |
| $d_N$ | 17.55*** | 2.82*** | 8.41*** | 0.46*** | 0.66*** | 29.89 |
| $d_S$ | 4.36*** | 0.44*** | 5.38*** | 2.16*** | 0.38*** | 12.72 |
| Percent contributions of each variable |  |  |  |  |  |  |
| Chaperone dependence | 0.10 | 0.05 | **0.71** | 0.08 | 0.06 |  |
| Number of protein–protein interactions | **0.42** | 0.02 | 0.00 | 0.02 | **0.54** |  |
| Expression level | **0.22** | 0.02 | **0.27** | **0.37** | 0.12 |  |
| Duplicability | 0.00 | **0.74** | 0.02 | 0.19 | 0.05 |  |
| Essentiality | **0.26** | 0.18 | 0.00 | **0.34** | **0.22** |  |

Significance levels: \* $P < 0.05$ , \*\* $P < 0.001$ , \*\*\* $P < 10^{-5}$ . We indicate in bold the contributions of a predictor to a component when greater than 20%.

**Fig. 3.**
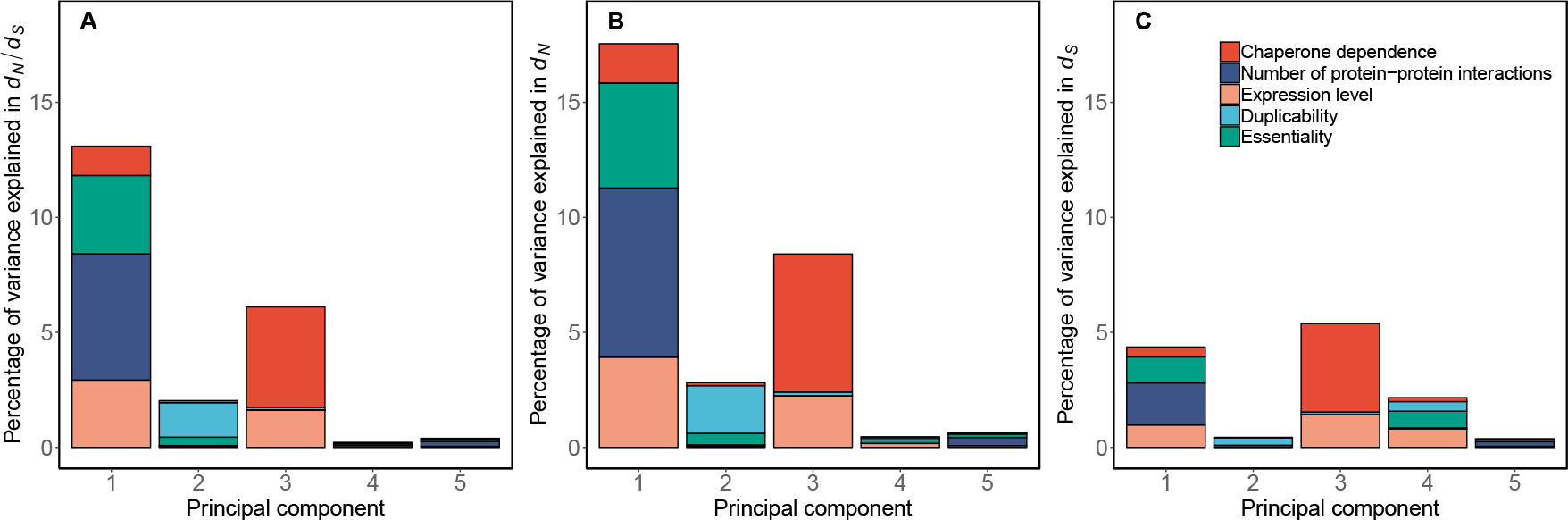
Principal component regression on (A) *d*_N_*/d*_S_, (B) *d*_N_, and (C) *d*_S_ in 5,532 yeast genes. For each principal component, the height of the bar represents the percent of variance in the rate of evolution explained by the component. The relative contribution of each variable to a principal component is represented with different colors. Table 3 contains the numerical data used to draw this figure.

For evolutionary rates measured as *d*_N_/*d*_S_, we found a principal component with a ~70% contribution of chaperone dependence and ~30% of expression level. This component explained a modest 6% of the variance with high significance (Table 2, Fig. 3A). Another significant principal component explains 13% of the variance. This component is mainly determined by the number of protein–protein interactions, essentiality, and expression level. A component explaining just ~2% of the variance was mainly determined by duplicability. The other two significant components explained in combination less than 1% of the variance. In summary, we found that chaperone dependence was the biological factor explaining the largest fraction of the total variance in the rate of evolution measured as *d*_N_/*d*_S_ (5.77%) (Table 4). It explained a larger fraction of the total variance than expression level (4.72%), and similar to the fraction explained by the number of protein–protein interactions (5.75%). Similar results were observed for *d*_N_ (Tables 3 and 4, Fig. 3). For *d*_S_, chaperone dependence was still the main factor explaining the total variance in the rate of evolution, with a contribution of 4.48% — still above that of expression level (3.27%) (Table 4). Indeed, it was the main determinant (~70%) of the principal component explaining the largest fraction of the variance (5.38%) (Table 3).

**Table 4.** Total variance explained by each variable in the principal component regression analysis of divergence data

| | $d_N/d_S$ | $d_N$ | $d_S$ |
| --- | --- | --- | --- |
| Chaperone dependence | 5.77% | 7.91% | 4.48% |
| Number of protein–protein interactions | 5.75% | 7.78% | 2.08% |
| Expression level | 4.72% | 6.47% | 3.27% |
| Duplicability | 1.68% | 2.36% | 0.86% |
| Essentiality | 3.93% | 5.37% | 2.03% |

Finally, we performed an analysis of covariance (ANCOVA), which is a category-based analysis in which we evaluated the effect of chaperone dependence on the rate of protein evolution measured as *d*_N_*/d*_S_ while controlling for the effect of the most important predictors: number of protein–protein interactions, expression level, and essentiality. We used the principal component of these three variables (principal component 1 in Table 3 and Fig. 3) as the continuous variable in the ANCOVA. We found that chaperone clients evolve on average 23% faster than all proteins (*P* = 8.6 × 10^−7^) (Fig. 4).

**Fig. 4.**
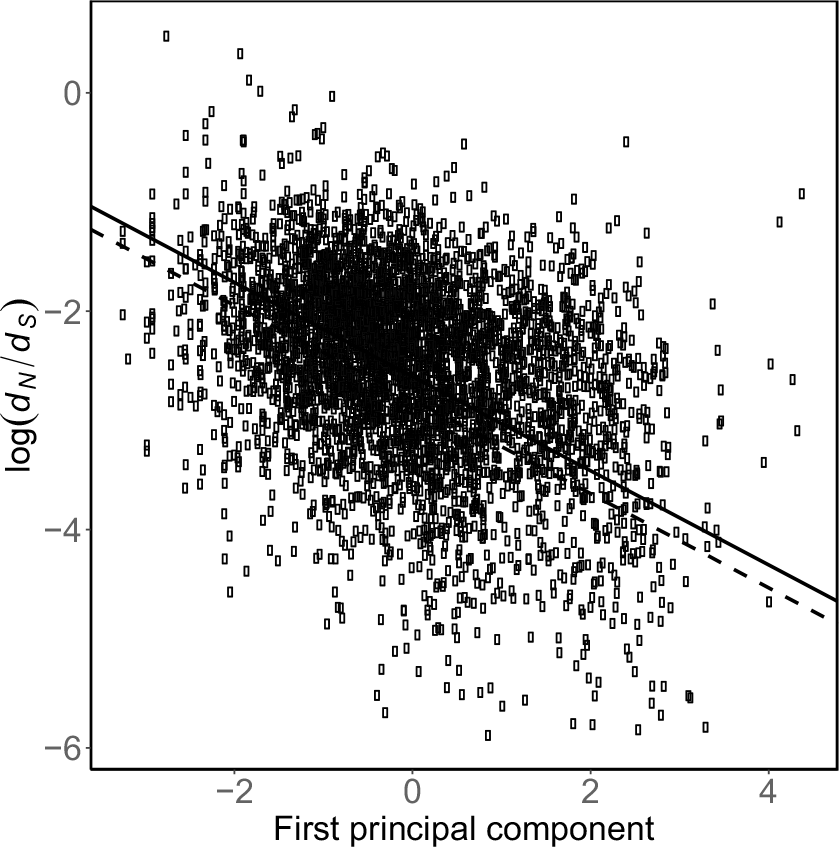
ANCOVA. Chaperone clients (gray points, continuous line) evolve 20% above the genome average rate (light points, dashed line) when considering divergence data between *S. cerevisiae* and *S. paradoxus*.

### Separate analysis of the clients of individual chaperones

Thus far we have aggregated the clients of all chaperones into a single group. However, different chaperones may affect the rates of protein evolution in different ways. We thus considered the clients of each chaperone separately. For each chaperone, we compared the clients of the chaperone vs. the proteins that are not clients of any chaperone. We again found that in all 35 cases clients exhibit a lower median and average *d*_N_/*d*_S_, with significant differences in 32 cases (Mann–Whitney *U* test, *P* < 0.05; Table 5). However, partial correlations between *d*_N_/*d*_S_ and chaperone dependence using degree as controlling variable were positive in 23 cases (significantly positive in 13 cases) and negative in 12 cases (significantly negative in 0 cases). This approach has the limitations that some chaperones have very few known clients, and that clients of the chaperone of interest may also be clients of other chaperones.

**Table 5.**
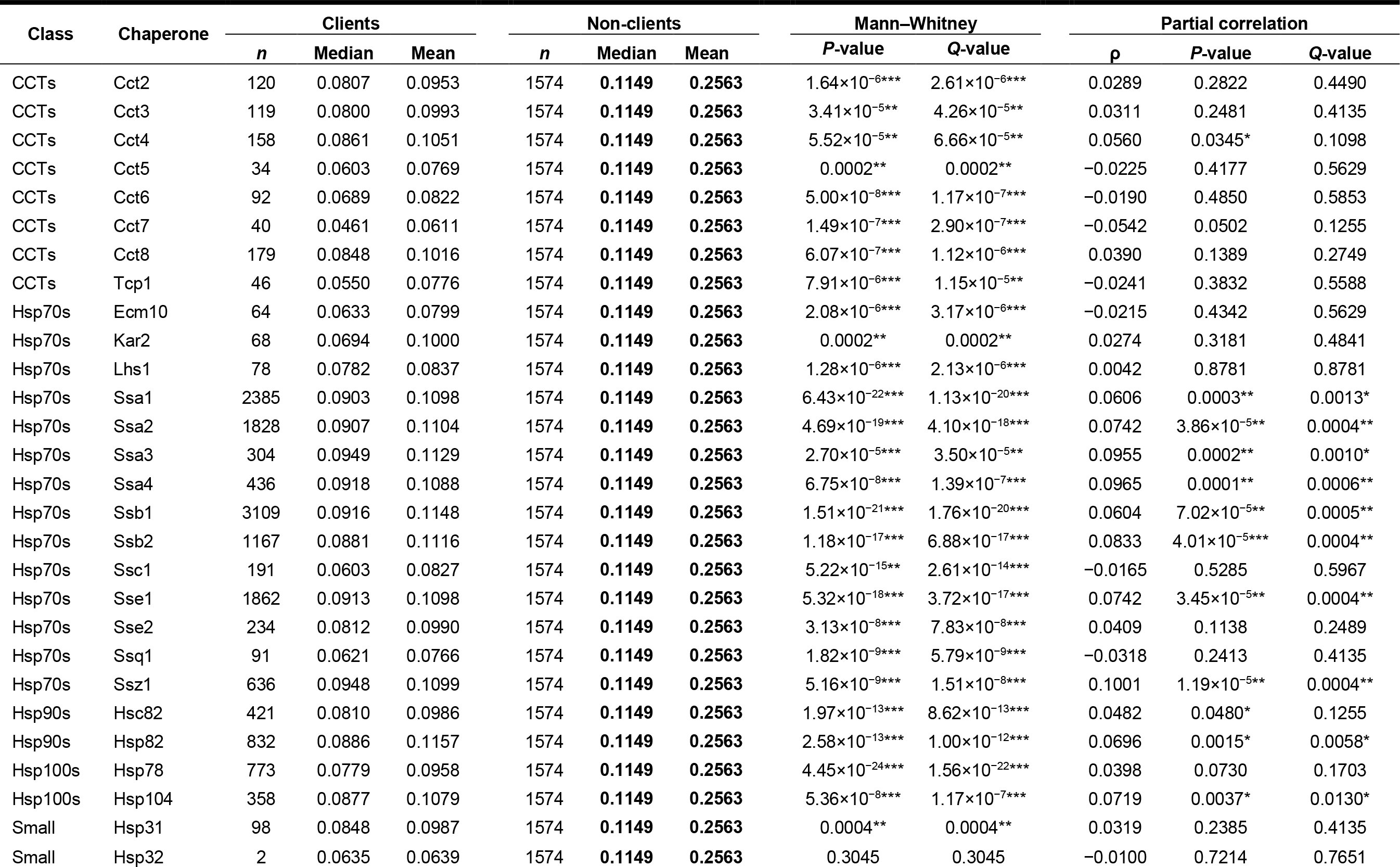
Rates of evolution of clients of different yeast chaperones vs. proteins that are not clients of any chaperone

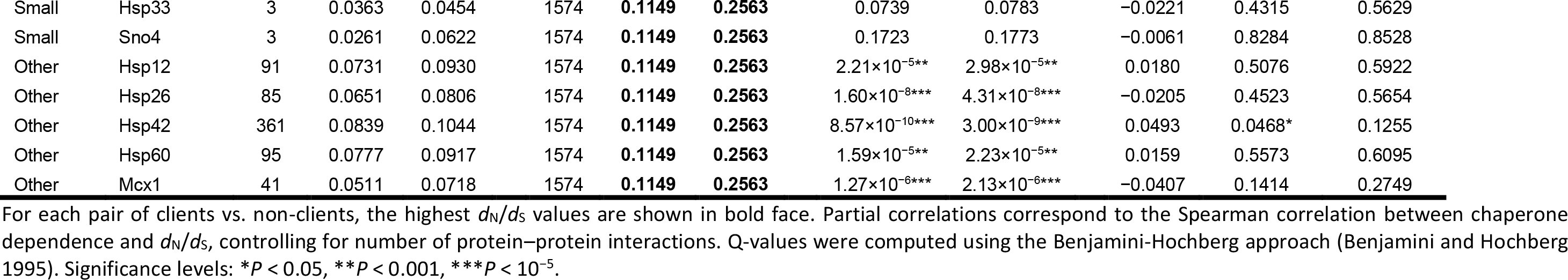

### Analysis of the clients of different groups of chaperones

We next grouped chaperones into five groups: small Hsps (Hsp31, Hsp32, Hsp33 and Sno4), Hsp70s (Kar2, Ssb1, Sse1, Sse2, Ssa1, Ssa2, Ssa3, Ssa4, Ssb2, Ecm10, Ssc1, Ssq1, Ssz1 and Lhs1), Hsp90s (Hsp82 and Hsc82), Hsp100s (Hsp78 and Hsp104) and CCTs (Tcp1, Cct4, Cct8, Cct2, Cct3, Cct5, Cct6, Cct7), and investigated the rates of evolution of the clients of each group. Single-family chaperones (Hsp26, Hsp42, Hsp12, Mcx1 and Hsp60) were not included in this analysis.

For each group of chaperones, we compared the rates of evolution of proteins that are clients of any of the chaperones of the group, vs. proteins that are not clients of any chaperone. In all 5 cases, clients had a significantly lower *d*_N_/*d*_S_. However, partial correlations between the dependence of each group and *d*_N_/*d*_S_ controlling for degree were always positive, and significant for the three chaperone classes with more clients (Hsp70s, Hsp90s and Hsp100s) (Table 6). This approach has the limitation that clients of one group of chaperones may also be clients of chaperones outside that group.

**Table 6.** Rates of evolution of clients of different chaperone groups vs. proteins that are not clients of any chaperone

| Class | Clients |  |  | Non-clients |  |  | Mann–Whitney |  | Partial correlation |  |  |
| --- | --- | --- | --- | --- | --- | --- | --- | --- | --- | --- | --- |
| | *n* | Median | Mean | *n* | Median | Mean | *P*-value | *Q*-value | $\rho$ | *P*-value | *Q*-value |
| CCTs | 614 | 0.0794 | 0.0967 | 1574 | **0.1149** | **0.2563** | $2.55 \times 10^{-20***}$ | $4.25 \times 10^{-20***}$ | 0.0349 | 0.1310 | 0.1638 |
| Hsp70s | 3783 | 0.0932 | 0.1156 | 1574 | **0.1149** | **0.2563** | $2.66 \times 10^{-21***}$ | $6.65 \times 10^{-21***}$ | 0.0550 | 0.0001** | 0.0005** |
| Hsp90s | 1101 | 0.0861 | 0.1115 | 1574 | **0.1149** | **0.2563** | $1.27 \times 10^{-17***}$ | $1.59 \times 10^{-17***}$ | 0.0615 | 0.0028* | 0.0070* |
| Hsp100s | 1004 | 0.0824 | 0.1005 | 1574 | **0.1149** | **0.2563** | $2.91 \times 10^{-23***}$ | $1.46 \times 10^{-22***}$ | 0.0537 | 0.0106* | 0.0176* |
| Small | 104 | 0.0809 | 0.0966 | 1574 | **0.1149** | **0.2563** | 0.0001** | 0.0001** | 0.0285 | 0.2926 | 0.2926 |

For each pair of clients vs. non-clients, the highest $d_N/d_S$ values are shown in bold face. Partial correlations correspond to the Spearman correlation between chaperone dependence and $d_N/d_S$ , controlling for number of protein–protein interactions. Q-values were computed using the Benjamini-Hochberg approach (Benjamini and Hochberg 1995). Significance levels: \* $P < 0.05$ , \*\* $P < 0.001$ , \*\*\* $P < 10^{-5}$ .

Next, in order to tease apart the effects of the different chaperone groups on rates of protein evolution while controlling for possible confounding factors, we performed two different multivariable analyses. We first performed a multiple linear regression analysis regressing *d*_N_*/d*_S_ against the four confounding biological factors we consider here (number of protein–protein interactions, expression level, essentiality, and duplicability), and dependence of the five chaperone families (Hsp70s, Hsp90s, Hsp100s, CTTs, and small Hsps). We found that among the chaperone families only Hsp70s and Hsp90s make a significant contribution to the regression and that the overall *R*^*2*^ is 0.220 (Table 7). Hsp70s and Hsp90s dependence were the only factors with a positive coefficient, indicating that dependence on these two major chaperone groups increases protein evolutionary rates. The contribution of Hsp90s was lost when regressing *d*_N_ or *d*_S_ instead of *d*_N_*/d*_S_ (Table 7).

**Table 7.** Multiple linear regression of different chaperone families

| | $d_N/d_S$ | $d_N$ | $d_S$ |
| --- | --- | --- | --- |
| HSP70 dependence | 0.15*** | 0.12*** | 0.04** |
| HSP90 dependence | 0.10* | 0.06 | 0.02 |
| HSP100 dependence | 0.04 | 0.05 | 0.05*** |
| CTT dependence | -0.03 | -0.04 | -0.01 |
| SMALL dependence | -0.10 | 0.02 | 0.03 |
| Number of protein–protein interactions | -0.16*** | -0.13*** | -0.01* |
| Expression level | -0.34*** | -0.31*** | -0.09*** |
| Duplicability | -0.38*** | -0.34*** | -0.02* |
| Essentiality | -0.35*** | -0.30*** | -0.01 |

Significance levels: \* $P < 0.05$ , \*\* $P < 0.001$ , \*\*\* $P < 10^{-5}$ .

We then performed a principal component regression analysis using the same predictor variables as above. Table 8 shows numerical data from the analysis, while Fig. 5 shows these data graphically. Neither Hsp70s dependence nor Hsp90s dependence contributed individually more than 20% to any significant principal component, but in combination they determine 30% of a component explaining 4.48% of the variance in *d*_N_*/d*_S_ (Table 8). In combination, Hsp70s and Hsp90s dependence contribute 3.19% to the total variance in the rate of evolution, which is above the contribution of the number of protein– protein interactions, but below the contributions of expression level, essentiality, or duplicability (Table 9).

**Table 8.**
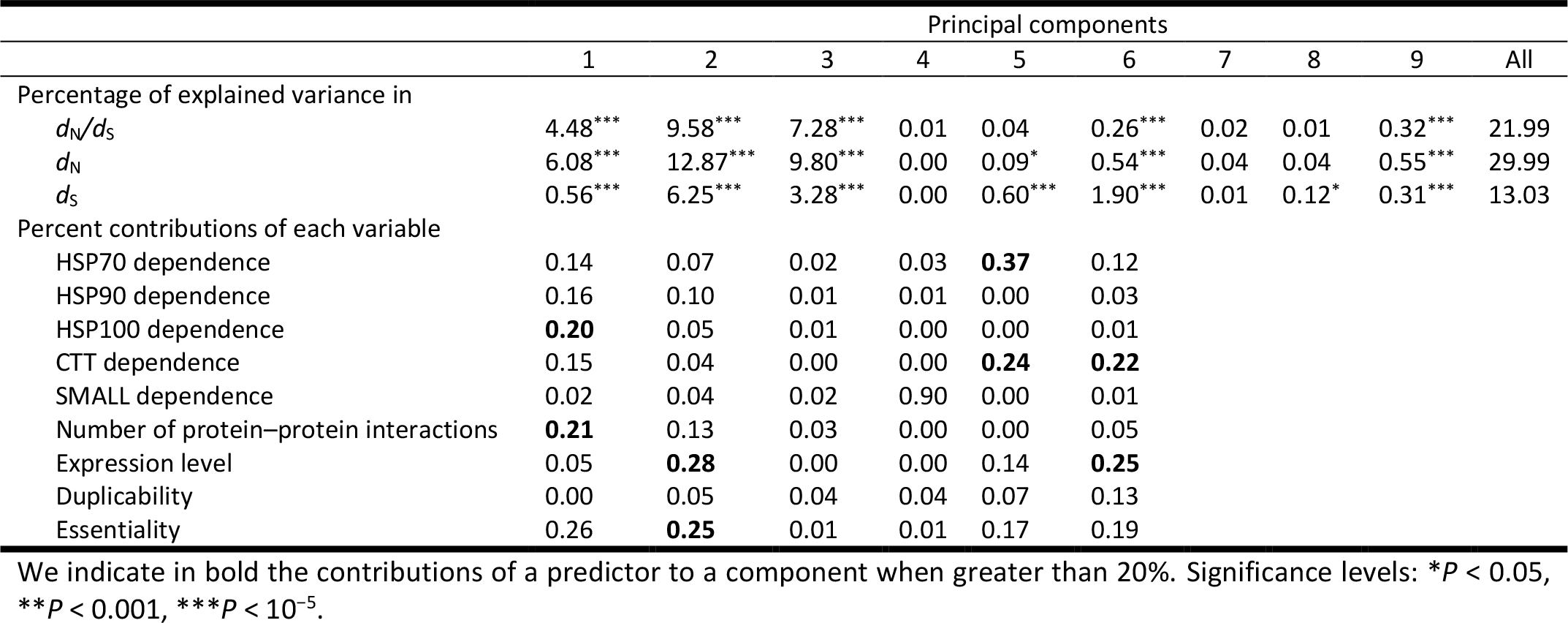
Results from the principal component regression analysis of divergence data

|  | Principal components |  |  |  |  |  |  |  |  |  |
| --- | --- | --- | --- | --- | --- | --- | --- | --- | --- | --- |
|  | 1 | 2 | 3 | 4 | 5 | 6 | 7 | 8 | 9 | All |
| Percentage of explained variance in |  |  |  |  |  |  |  |  |  |  |
| $d_N/d_S$ | 4.48*** | 9.58*** | 7.28*** | 0.01 | 0.04 | 0.26*** | 0.02 | 0.01 | 0.32*** | 21.99 |
| $d_N$ | 6.08*** | 12.87*** | 9.80*** | 0.00 | 0.09* | 0.54*** | 0.04 | 0.04 | 0.55*** | 29.99 |
| $d_S$ | 0.56*** | 6.25*** | 3.28*** | 0.00 | 0.60*** | 1.90*** | 0.01 | 0.12* | 0.31*** | 13.03 |
| Percent contributions of each variable |  |  |  |  |  |  |  |  |  |  |
| HSP70 dependence | 0.14 | 0.07 | 0.02 | 0.03 | **0.37** | 0.12 |  |  |  |  |
| HSP90 dependence | 0.16 | 0.10 | 0.01 | 0.01 | 0.00 | 0.03 |  |  |  |  |
| HSP100 dependence | **0.20** | 0.05 | 0.01 | 0.00 | 0.00 | 0.01 |  |  |  |  |
| CTT dependence | 0.15 | 0.04 | 0.00 | 0.00 | **0.24** | **0.22** |  |  |  |  |
| SMALL dependence | 0.02 | 0.04 | 0.02 | 0.90 | 0.00 | 0.01 |  |  |  |  |
| Number of protein–protein interactions | **0.21** | 0.13 | 0.03 | 0.00 | 0.00 | 0.05 |  |  |  |  |
| Expression level | 0.05 | **0.28** | 0.00 | 0.00 | 0.14 | **0.25** |  |  |  |  |
| Duplicability | 0.00 | 0.05 | 0.04 | 0.04 | 0.07 | 0.13 |  |  |  |  |
| Essentiality | 0.26 | **0.25** | 0.01 | 0.01 | 0.17 | 0.19 |  |  |  |  |

We indicate in bold the contributions of a predictor to a component when greater than 20%. Significance levels: \* $P < 0.05$ , \*\* $P < 0.001$ , \*\*\* $P < 10^{-5}$ .

**Table 9.** Total variance explained by each variable in the principal component regression analysis

| | $d_N/d_S$ | $d_N$ | $d_S$ |
| --- | --- | --- | --- |
| HSP70 dependence | 1.48% | 2.04% | 1.05% |
| HSP90 dependence | 1.71% | 2.32% | 0.80% |
| HSP100 dependence | 1.43% | 1.95% | 0.54% |
| CTT dependence | 1.10% | 1.54% | 0.88% |
| SMALL dependence | 0.64% | 0.84% | 0.35% |
| Number of protein–protein interactions | 2.66% | 3.66% | 1.35% |
| Expression level | 4.07% | 5.55% | 2.86% |
| Duplicability | 5.36% | 7.25% | 2.79% |
| Essentiality | 3.55% | 4.84% | 2.43% |

**Fig. 5.**
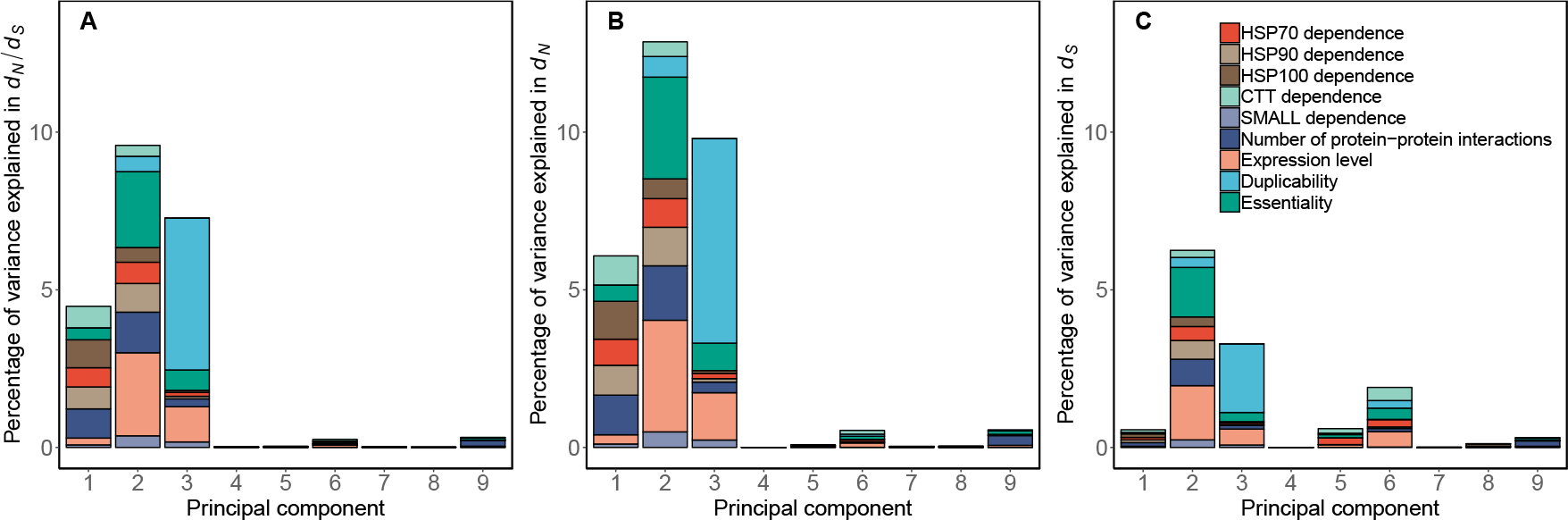
Principal component regression on (A) *d*_N_*/d*_S_, (B) *d*_N_, and (C) *d*_S_ in 5,532 yeast genes. For each principal component, the height of the bar represents the percent of variance in the rate of evolution explained by the component. The relative contribution of each variable to a principal component is represented with different colors. Table 10 contains the numerical data used to draw this figure.

### Chaperones increase the ratio of nonsynonymous to synonymous polymorphism ratio

For each *S. cerevisiae* gene, we obtained the nonsynonymous to synonymous polymorphism ratio (*d*_N_/*d*_S_) from Peter et al. (2018). Chaperone clients exhibit a significantly lower *d*_N_/*d*_S_ ratio (median for clients: 0.2352, median for non-clients: 0.2642, Mann–Whitney’s *U* test, *P* = 2.96×10^−10^). Partial correlation between *d*_N_/*d*_S_ and chaperone dependence controlling for expression level was nonsignificant (ρ = −0.0015, *P* = 0.9159), and the partial correlation between *d*_N_/*d*_S_ and chaperone dependence controlling for network degree was significantly positive (ρ = 0.0612, *P* = 10^−5^), indicating that chaperones accelerate rates of protein evolution.

**Table 10.**
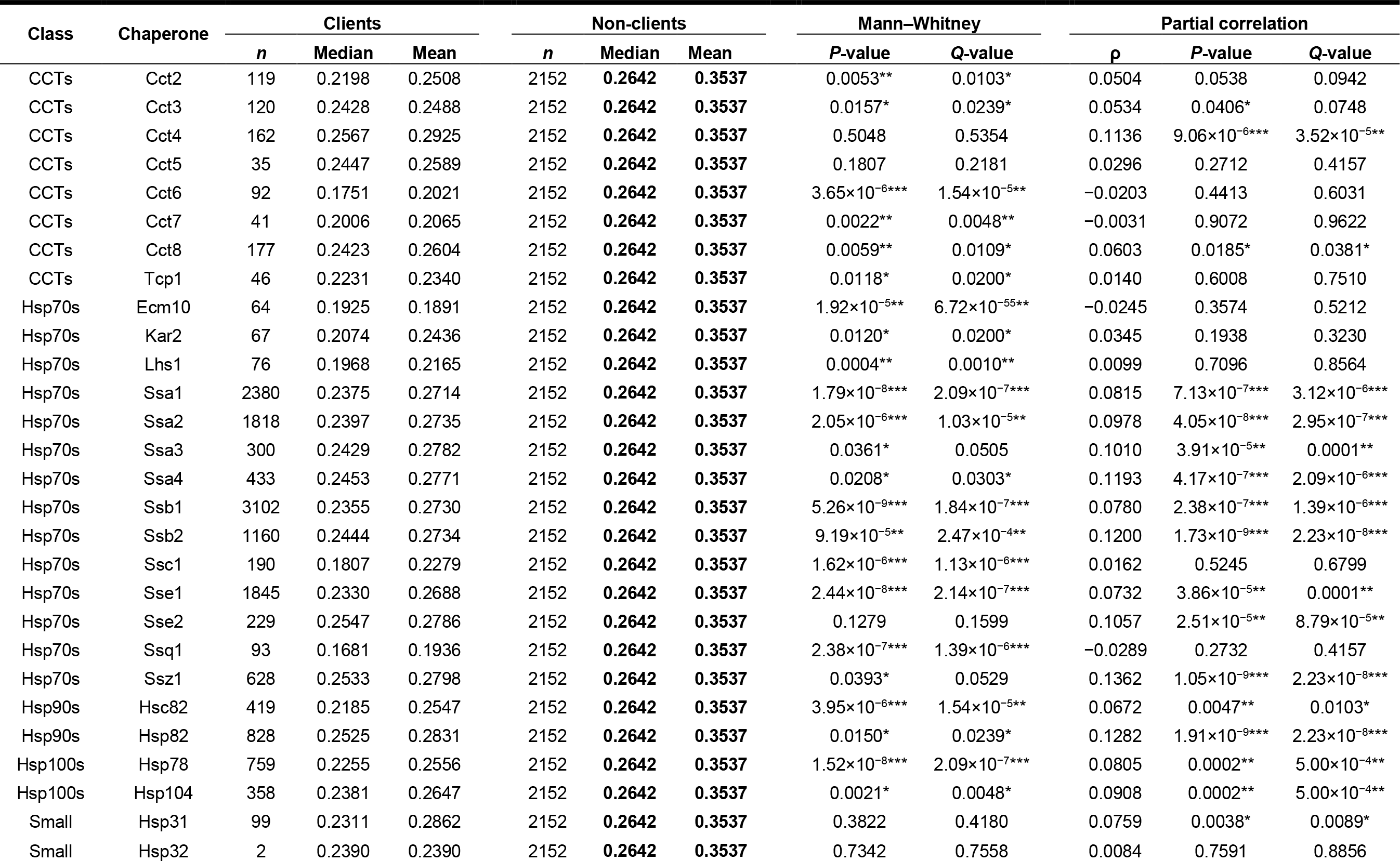
Nonsynonymous to synonymous polymorphism ratio of clients of different yeast chaperones vs. proteins that are not clients of any chaperone

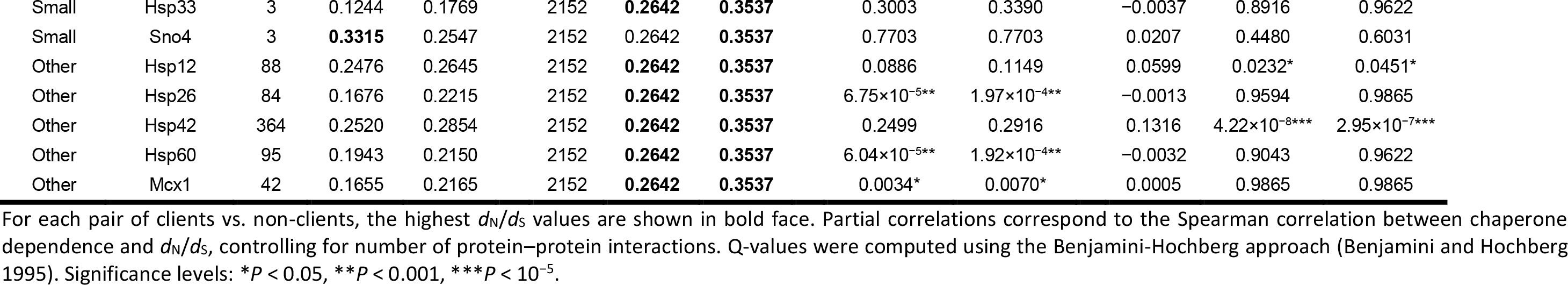

For each chaperone, we compared the rates of evolution of their clients (*n* ranged from 2 to 3102) vs. the rates of evolution of non-clients (proteins that are not clients of any chaperone, *n* = 2152). In all 35 cases, clients exhibited a lower average *d*_N_/*d*_S_, and in 34 of the cases they also exhibited a lower median *d*_N_/*d*_S_, with significant differences in 26 cases (Mann– Whitney’s *U* test, *P* < 0.05; Table 10). Partial correlations between *d*_N_/*d*_S_ and chaperone dependence controlling for degree were positive in 28 cases (significantly positive in 19 cases) and negative in 7 cases (significantly negative in 0 cases).

Next, for each group of chaperones (small Hsps, Hsp70s, Hsp90s, Hsp100s and CCTs), we compared the rates of non-synonymous to synonymous polymorphism of the clients of any of the group (*n* ranged from 103 to 947) vs. those of proteins that are not clients of any chaperone (*n* = 2152). In all 5 cases clients exhibited lower median and mean *d*_N_/*d*_S_, with significant differences (Mann–Whitney’s *U* test, *P* < 0.05) in all cases except for the clients of small Hsps (the smallest group; Table 11). However, partial correlations between *d*_N_/*d*_S_ and chaperone dependence controlling for network degree was always significantly positive.

**Table 11.** Nonsynonymous to synonymous polymorphism ratio of clients of different chaperone groups vs. proteins that are not clients of any chaperone

| Class | Clients |  |  | Non-clients |  |  | Mann-Whitney |  | Partial correlation |  |  |
| --- | --- | --- | --- | --- | --- | --- | --- | --- | --- | --- | --- |
| | *n* | Median | Mean | *n* | Median | Mean | *P*-value | *Q*-value | $\rho$ | *P*-value | *Q*-value |
| CCTs | 487 | 0.2436 | 0.2694 | 2152 | **0.2642** | **0.3537** | 0.0004** | 0.0001** | 0.0942 | 5.36×10<sup>-5</sup>** | 0.0020* |
| Hsp70s | 836 | 0.2416 | 0.2907 | 2152 | **0.2642** | **0.3537** | 0.0009** | 0.0201* | 0.0504 | 0.0201* | 0.0352* |
| Hsp90s | 947 | 0.2433 | 0.2731 | 2152 | **0.2642** | **0.3537** | 0.0001** | 4.60×10<sup>-6</sup>*** | 0.1024 | 9.20×10<sup>-7</sup>*** | 0.0003** |
| Hsp100s | 863 | 0.2272 | 0.2592 | 2152 | **0.2642** | **0.3537** | 1.82×10<sup>-8</sup>*** | 0.0007** | 0.0752 | 0.0004** | 0.0120* |
| Small | 103 | 0.2479 | 0.2875 | 2152 | **0.2642** | **0.3537** | 0.4503 | 0.0020* | 0.0825 | 0.0016* | 0.0066* |

For each pair of clients vs. non-clients, the highest $d_N/d_S$ values are shown in bold face. Partial correlations correspond to the Spearman correlation between chaperone dependence and $d_N/d_S$ , controlling for number of protein–protein interactions. *Q*-values were computed using the Benjamini-Hochberg approach (Benjamini and Hochberg 1995). Significance levels: \* $P < 0.05$ , \*\* $P < 0.001$ , \*\*\* $P < 10^{-5}$ .

Finally, we performed a multivariable analysis to study the effect of chaperone dependence on *d*_N_/*d*_S_ at the intra-population level controlling simultaneously for all the studied variables, as we did previously for the divergence data. The results are very similar. We first regressed *d*_N_*/d*_S_ against the five biological factors, and found that all make a significant contribution to the regression and that the overall *R*^*2*^ is 0.17 (Table 12). Chaperone dependence was the only factor with a positive coefficient, indicating that chaperone dependence also increases *d*_N_/*d*_S_ within yeast populations. We also performed a principal component regression analysis using the same predictor variables as above. Table 13 shows numerical data from the analysis, while Fig. 6 shows these data graphically.

**Table 12.** Multiple linear regression

| | $d_N/d_S$ |
| --- | --- |
| Chaperone dependence | 0.16<sup>***</sup> |
| Number of protein–protein interactions | −0.06<sup>***</sup> |
| Expression level | −0.23<sup>***</sup> |
| Duplicability | −0.06<sup>*</sup> |
| Essentiality | −0.20<sup>***</sup> |

Significance levels: $^*P < 0.05$ , $^{**}P < 0.001$ , $^{***}P < 10^{-5}$ .

**Table 13.**
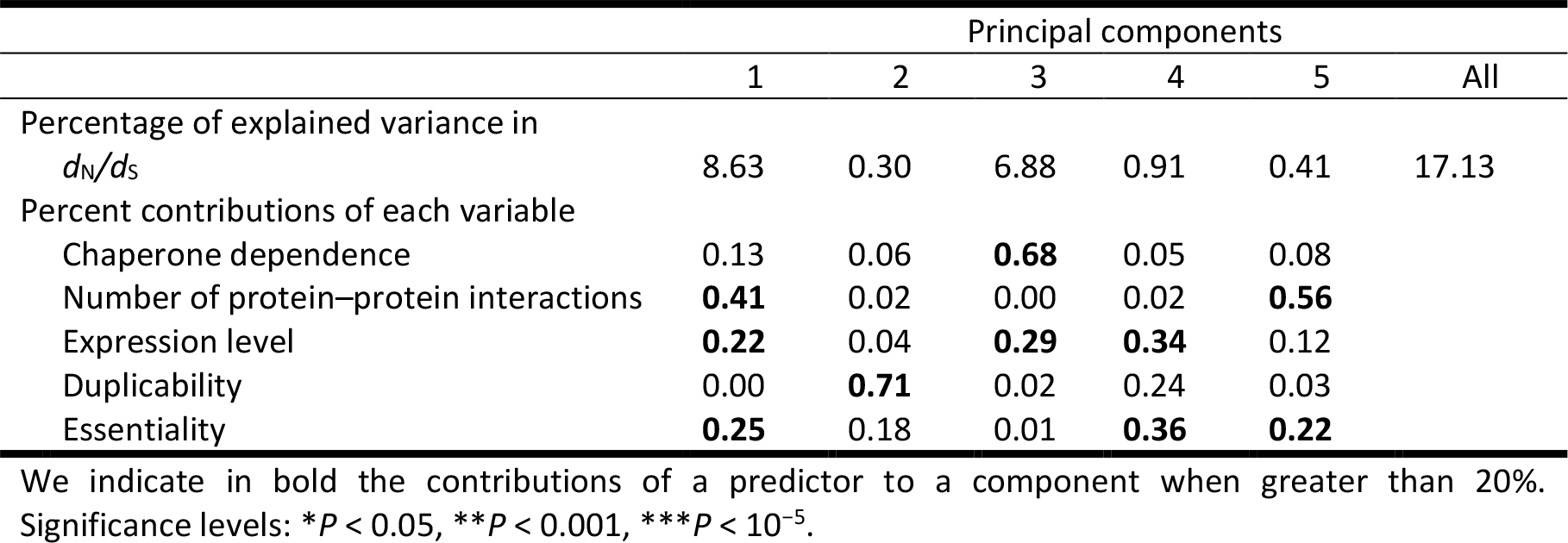
Results from the principal component regression analysis of polymorphism data

|  | Principal components |  |  |  |  |  |
| --- | --- | --- | --- | --- | --- | --- |
|  | 1 | 2 | 3 | 4 | 5 | All |
| Percentage of explained variance in $d_N/d_S$ | 8.63 | 0.30 | 6.88 | 0.91 | 0.41 | 17.13 |
| Percent contributions of each variable |  |  |  |  |  |  |
| Chaperone dependence | 0.13 | 0.06 | **0.68** | 0.05 | 0.08 |  |
| Number of protein–protein interactions | **0.41** | 0.02 | 0.00 | 0.02 | **0.56** |  |
| Expression level | **0.22** | 0.04 | **0.29** | **0.34** | 0.12 |  |
| Duplicability | 0.00 | **0.71** | 0.02 | 0.24 | 0.03 |  |
| Essentiality | **0.25** | 0.18 | 0.01 | **0.36** | **0.22** |  |

We indicate in bold the contributions of a predictor to a component when greater than 20%. Significance levels: \* $P < 0.05$ , \*\* $P < 0.001$ , \*\*\* $P < 10^{-5}$ .

**Fig. 6.**
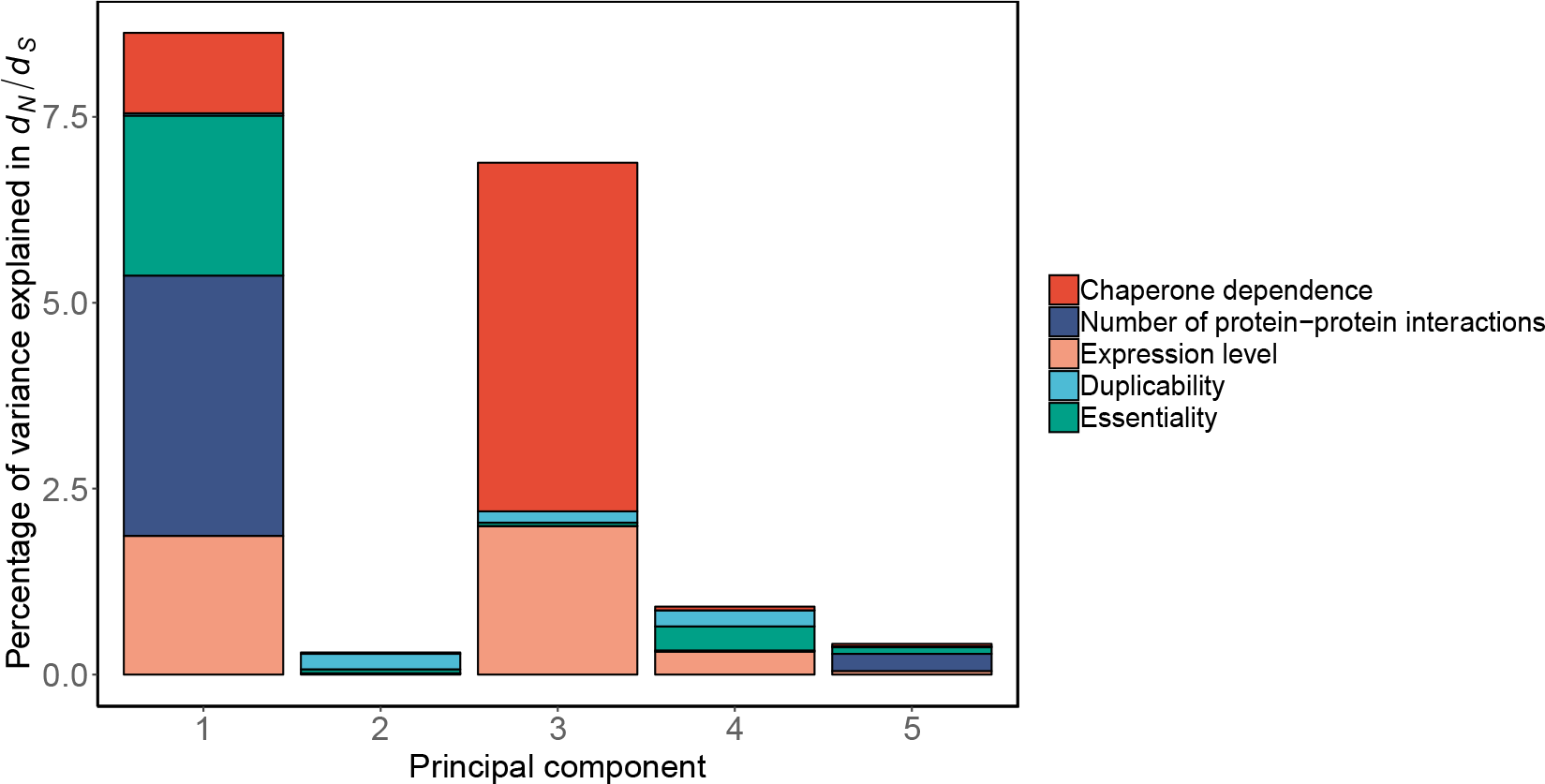
Principal component regression on *d*_N_*/d*_S_ in 6,132 yeast genes. For each principal component, the height of the bar represents the percent of variance in the rate of evolution explained by the component. The relative contribution of each variable to a principal component is represented with different colors. Table 13 contains the numerical data used to draw this figure.

As with divergence data, we found a principal component with a 70% contribution of chaperone dependence and 30% expression level. This component explained ~7% of the variance of *d*_N_/*d*_S_ (Table 13, Fig. 6). Another significant principal component explains 8.6% of the variance. This component is mainly determined by the number of protein–protein interactions, essentiality, and expression level. The other three significant components explained in combination less than 2% of the variance. In summary, we also found that chaperone dependence was the biological factor explaining the largest fraction of the total variance in the rate of evolution measured as *d*_N_/*d*_S_ (5.87%), explaining a larger fraction of the total variance than expression level (4.23%) and the number of protein–protein interactions (3.76%) (Table 14).

**Table 14.** Total variance explained by each variable in the principal component regression analysis

| | $d_N/d_S$ |
| --- | --- |
| Chaperone dependence | 5.87% |
| Number of protein–protein interactions | 3.76% |
| Expression level | 4.23% |
| Duplicability | 0.62% |
| Essentiality | 2.66% |

Finally, we performed an ANCOVA to evaluate the effect of chaperone dependence on the rate of protein evolution while controlling for the effect of the number of protein–protein interactions, expression level, and essentiality. As the continuous variable in the ANCOVA, we used the principal component of these three variables (principal component 1 in Table 13 and Fig. 6). We found that chaperone clients evolve on average 19.2% faster than the proteome average (*P* = 3.6 × 10^−11^) (Fig. 7).

**Fig. 7.**
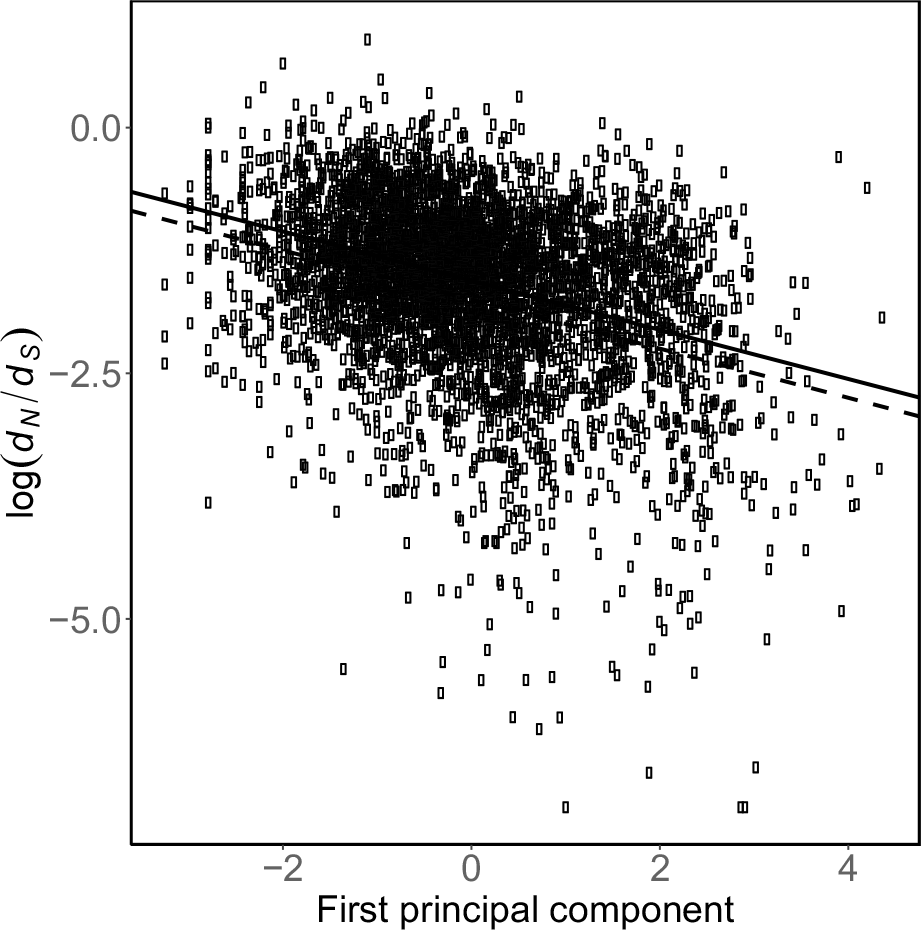
ANCOVA. Chaperone clients (gray points, continuous line) evolve 19.2% above the genome average rate (light points, dashed line) when considering genetic variants segregating in *S. cerevisiae*.

## DISCUSSION

We study how the different yeast chaperones affect the evolutionary rate of their protein clients. In particular, we analyze the effect of chaperone dependence on protein evolution at two very different evolutionary time scales. We first study how chaperone-mediated folding has affected protein evolution over the evolutionary divergence of *S. cerevisiae* and *S. paradoxus*. We then study if the same process has left a signature on the patterns of standing genetic variation found in modern wild and domesticated strains of *S. cerevisiae* (Peter, et al. 2018). We find that chaperone-mediated buffering has indeed left a trace on the protein-coding regions of the yeast genome, such that genes encoding chaperone clients (“client genes”) have diverged faster than genes encoding nonclient proteins (“nonclient genes”) when controlling for their number of protein-protein interactions. We also find that client genes have accumulated more genetic diversity than nonclients genes among natural strains of *S. cerevisiae.* In a principal component regression analysis, we find that chaperone dependence explains the largest fraction of the observed variance in the rate of evolution at both evolutionary time scales. This contribution of chaperone-mediated folding to the variations on the rate of protein evolution is well above the fraction of the variance explained by other well-known factors that affect protein evolution such as expression level or protein-protein interactions (Pál, et al. 2001; Fraser, et al. 2002; Drummond, et al. 2005).

Cost-benefit trade-offs are common in evolution, including protein evolution. Proteins are marginally stable (DePristo, et al. 2005) and soluble (Tartaglia, et al. 2007) inside a cell and their native structure is sensitive to mutations. Protein stability is a major constraint on protein evolution (Bloom, et al. 2006; Zeldovich, et al. 2007). Most nonsynonymous mutations diminish protein stability or solubility, and are therefore deleterious (Dobson 1999). Moreover, neofunctionalizing mutations that confer new protein functions, including new protein-protein interactions, tend to be highly destabilizing (Tokuriki, et al. 2008; Soskine and Tawfik 2010). Therefore, in the absence of chaperone buffering, the cost of a neofunctionalizing mutation may be larger than its benefit (Tokuriki and Tawfik 2009). Chaperones, by diminishing the negative effect of mutations on protein stability and folding, can promote protein evolution, and potentiate the regulatory or metabolic effect of a protein mutation (Taipale, et al. 2010). Our finding that yeast chaperones can accelerate protein evolution is in line with previous observations that chaperones can act as evolutionary capacitors (Queitsch, et al. 2002; Rutherford 2003; Jarosz and Lindquist 2010), buffer the destabilizing effect of mutations (Tokuriki and Tawfik 2009), facilitate the divergence of gene duplicates (Lachowiec, et al. 2013), and ultimately allow proteins to explore a larger fraction of their sequence space (Williams and Fares 2010; Pechmann and Frydman 2014; Aguilar-Rodríguez, et al. 2016; Kadibelban, et al. 2016). However, it is important to notice that chaperones do not just modify the effects of mutations affecting protein stability or folding. While most mutation are likely unaffected by chaperone activity, they can also modify (either buffer or potentiate) the fitness or phenotypic effects of mutations indirectly since they tend to be highly epistatic hubs whose influence can percolate through protein-protein interaction or regulatory networks (Taipale, et al. 2010; Zabinsky, et al. 2018). Even if chaperone-buffered genetic variants are only rarely acquired, they could be enriched in a population if stabilizing selection does not remove them (because their deleterious phenotypic consequences are masked by a chaperone). A recent study has found evidence for this hypothesis among Hsp90-dependent variants in *S. cerevisiae* that affect cell size and shape (Geiler-Samerotte, et al. 2016). We also find evidence for this enrichment of cryptic genetic variants within client genes among genetic variants segregating in *S. cerevisiae.*

Nonsynonymous mutations that allow the establishment of new physical interactions with other proteins are a class of neofunctionalizing mutations that can be highly destabilizing (Pechmann and Frydman 2014). Therefore, some chaperones can also buffer mutations that re-wire protein interactions, thus promoting the evolution of protein networks (Pechmann and Frydman 2014), and perhaps explaining why chaperone clients tend to be well-connected in such networks, as we observed here. Furthermore, chaperones and their clients coevolve in a process where sequence changes in the chaperone may lead to compensatory changes in their clients and further re-wiring of the protein networks they form (Koubkova-Yu, et al. 2018).

In a multivariable statistical analysis, we find that the chaperones affecting rates of protein evolution belong to two major chaperone families: Hsp70s and Hsp90s. While there is ample evidence that Hsp90s can accelerate the rate of protein evolution in other eukaryotic species (Lachowiec et al. 2013, Pechmann and Frydman 2014, Lachowiec et al. 2015), the evidence for eukaryotic Hsp70 chaperones having a similar effect is not so abundant. A previous study found that the ribosome-associated Hsp70 SSB chaperone that preferentially binds long and disordered nascent polypeptide chains accelerates the rate of accumulation of mutations likely to be destabilizing among weakly-interacting clients (Pechmann and Frydman 2014). In a previous study, we found that bacterial DnaK, which belongs to the same major chaperone family, also accelerates protein evolution using a combination of experimental and comparative genomics approaches (Aguilar-Rodríguez, et al. 2016). While it has been shown before that the chaperonin GroEL accelerates protein evolution (Bogumil and Dagan 2010; Williams and Fares 2010), we do not find good evidence here that the eukaryotic chaperonin system CCT, present in eukarya and archaea but absent from bacteria, has the same effect on protein evolution. We find that the chaperone Hsp104 from the Hsp100 family accelerates the evolution of its protein clients when controlling for number of protein-protein interactions. This could be the first observation that this important chaperone could affect protein evolutionary rates. However, we do not observe any effect of the family Hsp100 (Hsp78 and Hsp104) when controlling for possible confounding variables in a multiple linear regression and in a principal component regression analysis. Finally, we do not detect any significant effect of small heat shock proteins in the rate of evolution of their clients.

In summary, we analyzed the evolution of proteins that are subjected to folding assisted by different chaperones in the complex yeast chaperone network over two different evolutionary time scales. Our comparative approach indicates that chaperone-assisted folding increases the rate of protein evolution when properly controlling for confounding factors at both time scales. We show how protein chaperones, by virtue of their role in modulating protein genotype-phenotype maps, have a disproportionate effect on the evolution of the protein-coding regions of a genome. Our results highlight the importance of integrating different cellular factors when studying protein sequence evolution.

## METHODS

### Rates of protein evolution

The *S. cerevisiae* and *S. paradoxus* protein and coding (CDS) sequences were obtained from the *Saccharomyces* Genome Database (Cherry, et al. 2012). Each *S. cerevisiae* protein sequence was used as query in a BLASTP search (*E*-value cutoff = 10^−10^) against the *S. paradoxus* proteome. Similarly, each *S. paradoxus* protein was used in a BLASTP search against the *S. cerevisiae* proteome. Pairs of best reciprocal hits were considered to be encoded by pairs of orthologs. For each pair of orthologs, protein sequences were aligned using ProbCons (Do, et al. 2005), and the resulting alignments were used to guide the alignment of the corresponding CDSs. PAML version 4.4d (codeml program, M0 model; Yang 2007) was used to estimate *d*_N_, *d*_S_ and *d*N/*d*S values.

### Positive selection analyses

The *S. cerevisiae*, *S. paradoxus*, *S. mikatae*, *S. kudriavzevii*, and *S. bayanus* protein and CDS sequences were obtained from the *Saccharomyces* Genome Database (Cherry, et al. 2012). Each *S. cerevisiae* protein sequence was used as query in a BLASTP search (*E*-value cutoff = 10^−10^) against the *S. paradoxus*, *S. mikatae*, *S. kudriavzevii*, and *S. bayanus* proteomes. Similarly, each *S. paradoxus*, *S. mikatae*, *S. kudriavzevii*, and *S. bayanus* protein was used in a BLASTP search against the *S. cerevisiae* proteome. Pairs of best reciprocal hits were considered to be encoded by pairs of orthologs. Only genes with putative orthologs in *S. paradoxus*, *S. mikatae*, *S. kudriavzevii*, and *S. bayanus* were retained for analysis. For each groups of orthologs, protein sequences were aligned using ProbCons (Do, et al. 2005), and the resulting alignments were used to guide the alignment of the corresponding CDSs. Alignments were filtered as in (Luisi, et al. 2015).

The filtered alignments were used in tests of positive selection using PAML version 4.4d (codeml program, M8 vs. M7 test; Yang, et al. 2000). Twice the difference in the log-likelihood of both models was assumed to follow a χ^2^ distribution with two degrees of freedom. Genes with a *P*-value lower than 0.05 and a fraction of codons with *d*_N_/*d*_S_ higher than 1 were assumed to be under positive selection. All computations were run using three starting *d*_N_/*d*_S_ values (0.04, 0.4 and 4) in order to alleviate the problem of local optima. The alignments corresponding to genes with signatures of positive selection were visualized using BioEdit version 7.2.5 in order to discard alignment or annotation errors.

### Chaperone client data

Chaperone–client interaction data were obtained from Gong et al. (2009). Their study included 35 chaperones and 29 co-chaperones. For each chaperone, we obtained a list of clients from their supplementary table 2.

### Additional information

For each *S. cerevisiae* gene, the following information was gathered from different sources. The nonsynonymous to synonymous polymorphism ratio was obtained from Peter et al. (2018)). For each gene, the average *d*_N_/*d*_S_ across all pairs of genomes was used. Gene expression (mRNA abundance) levels were obtained from Nagalakshmi et al. (2008). The number of protein–protein interactions (degree centrality) was obtained from the BioGRID database, version v3.2.101. Only physical, non-redundant interactions among *S. cerevisiae* proteins were included in the analysis. Degrees were recomputed on a high-quality subnetwork, including those interactions determined by low-throughput studies or by more than one high-throughput study. A list of paralogs was obtained from Ensembl’s Biomart (Kinsella, et al. 2011), and genes with at least one paralog were classified as duplicates. A list of essential genes was obtained from Giaever et al. (2002).

### Statistical analyses

Statistical analyses were conducted using the R package (R Core Team 2014). Partial correlation analyses were conducted using the ‘pcor.test’ function (Kim 2015). We used the package ‘pls’ to carry out the principal component regression analysis. We carried out base-10 logarithmic transformations of the continuous variables when such transformations led to a higher *R*^*2*^. If a continuous variable contained values equal to zero, we added a small constant (0.001) to all its values to allow its logarithmic transformation. We scaled the independent variables to zero mean and unit variance.

## ACKNOWLEDGEMENTS

D.A.-P. and J.A.-R. dedicate this manuscript to the memory of M.A.F. D.A.-P. was supported by a grant from the National Science Foundation (MCB 1818288), funds from the University of Nevada, Reno, pilot grants from Nevada INBRE (P20GM103440) and the Smooth Muscle Plasticity COBRE from the University of Nevada, Reno (5P30GM110767-04), both funded by the National Institute of General Medical Sciences (National Institutes of Health), and a Juan de la Cierva postdoctoral fellowship from the Spanish Ministerio de Economía y Competitividad, Spain (JCA-2012-14056). J.A.-R. acknowledges support by Swiss National Science Foundation grant P2ZHP3_174735. M.A.F. was supported by grants from Science Foundation Ireland (12/IP/1637) and the Spanish Ministerio de Economía y Competitividad, Spain (MINECO-FEDER; BFU201236346 and BFU2015-66073-P).

